# WolfPackR: An R package for identifying wolf packs based on genetic and spatial data

**DOI:** 10.64898/2026.04.28.721440

**Authors:** Etienne Boncourt

## Abstract

The global expansion of grey wolf (*Canis lupus*) populations, particularly in Europe, underscores the need for robust tools to study their social structure, territory use, and genetic relatedness. Wolf packs are dynamic, evolving through dispersal, mortality, and reproductive success, and their accurate identification is crucial for effective conservation and conflict mitigation. Traditional methods for estimating wolf populations and pack structures—such as snow tracking or howling surveys—are labor-intensive and often unreliable. Noninvasive genetic sampling and spatial capture-recapture models have improved monitoring, but integrating genetic and spatial data remains a challenge.

We introduce WolfPackR, an R package designed to integrate genetic relatedness and spatial data for identifying wolf packs, lone individuals, and spatially isolated but genetically linked “ugly ducklings.” WolfPackR uses pairwise relatedness estimators to define genetic groups and refines these groups through spatial overlap analysis based on Minimum Convex Polygons (MCPs). The package provides a comprehensive toolkit for analyzing population structure, territoriality, and social organization, including functions for genetic grouping, spatial clustering, summary statistics, and interactive visualization.

We demonstrate the utility of WolfPackR using a case study of 505 genotyped and geospatialized wolf scat samples from Romania. By combining genetic and spatial data, WolfPackR accurately identifies pack structures that align with expert assessments and family tree reconstructions. The package modular design and reliance on widely used R libraries (dplyr, igraph, sf, leaflet) ensure flexibility and ease of integration into existing workflows. While sampling heterogeneity may limit territory delineation in some cases, WolfPackR offers a cost-effective and reproducible framework for studying wolf pack dynamics, with potential applications for other social species.

## Introduction

Grey wolf (*Canis lupus*) populations are expanding across multiple regions globally, particularly in Europe, where they are recolonizing areas following historical declines in the 20th century (Chapron et al., 2014). This resurgence is a testament to successful conservation efforts and changing societal attitudes toward large carnivores (Boitani, 2003). Wolves are highly social animals that live in structured family groups known as packs, typically consisting of a breeding pair and their offspring from previous years (Mech & Boitani, 2003). Packs occupy and defend territories, which can vary in size depending on prey availability, human disturbance, and landscape features (Fuller et al., 2003). The social structure of wolf packs is complex, involving not only genetic relationships between parents and offspring but also the integration of unrelated individuals, such as immigrants or dispersers (Iosif et al., 2025; Mech & Boitani, 2003; Rutledge et al., 2010). These dynamics are crucial for maintaining genetic diversity and ensuring the long-term viability of wolf populations (Borg et al., 2015).

Understanding wolf pack structure and territory use is essential for effective management and conservation. Wolf packs are not static; they evolve over time due to dispersal, mortality, and reproductive success (Ausband et al., 2017). For example, the loss of a breeding individual can lead to pack dissolution or the integration of new members, which can affect social cohesion and territory stability (Borg et al., 2015). Additionally, wolves may form complex social relationships with unrelated individuals, who can contribute to pack activities and territory defense (Pacheco et al., 2024). These dynamics highlight the importance of considering both genetic and social factors when studying wolf populations.

Wolf management is a contentious issue, particularly in regions where wolves coexist with human activities such as livestock farming (Treves & Karanth, 2003). Effective management strategies require accurate population estimates and an understanding of pack dynamics to mitigate conflicts and promote coexistence (Redpath et al., 2017). Traditional methods for estimating wolf populations, such as snow tracking or howling surveys, are labor-intensive and can be unreliable, especially in areas with low snow cover or dense vegetation (Alexander et al., 2005). In recent decades, noninvasive genetic sampling has emerged as a powerful tool for monitoring wolf populations (Schwartz et al., 2007). This approach involves collecting DNA from sources such as scat, hair, or urine, which are deposited by wolves in their environment (Kelly et al., 2012). Noninvasive sampling is particularly advantageous because it minimizes disturbance to the animals and can be conducted over large and remote areas (Taberlet, 1996).

The advent of noninvasive DNA sampling has revolutionized wildlife monitoring by providing a cost-effective and scalable method for identifying individuals, reconstructing pedigrees, and estimating population parameters (Caniglia et al., 2014). For example, genetic data can be used to infer relatedness among individuals, identify parent-offspring relationships, and detect hybridization events with domestic dogs (Pilot et al., 2021). When combined with spatial data, such as GPS locations or territory mapping, genetic information can provide a comprehensive understanding of pack structure and dynamics (López-Bao et al., 2018). For instance, spatial capture-recapture models, which integrate spatial and genetic data, have become increasingly popular for estimating population density and assessing spatial patterns of animal distributions (Royle et al., 2014).

Despite these advancements, there are still challenges in accurately identifying wolf packs and their territories. Traditional genetic analysis tools, such as COLONY (Jones & Wang, 2010) or STRUCTURE (Pritchard et al., 2000), focus primarily on relatedness and may not account for spatial dynamics, leading to potential misclassification of individuals that migrate or are spatially isolated (Manel et al., 2003). Conversely, spatial clustering tools often ignore genetic context, which can result in incomplete or biased pack assignments (Calenge, 2006). To address these limitations, integrated approaches that combine genetic and spatial data are needed to improve the accuracy of pack identification and territory delineation (Iosif et al., 2025).

In this context, `WolfPackR` offers a novel solution by integrating genetic relatedness and spatial data to identify wolf packs, lone individuals, and “ugly ducklings” (genetically linked but spatially isolated individuals). The package uses pairwise relatedness estimators, such as Wang’s estimator (Wang, 2002), to define potential genetic groups and refines these groups using spatial overlap analysis based on Minimum Convex Polygons (MCPs). This approach provides a more accurate classification of individuals into packs, accounting for both genetic and spatial dynamics. `WolfPackR` is designed for researchers working with genetic relatedness matrices (e.g., from microsatellite or SNP data) and spatial location data (e.g., from scat locations). By combining genetic networks (using `igraph`; Csárdi et al., 2025), spatial overlap analysis (computing MCPs with `sf`; Pebesma, 2018), and visualization tools (using `leaflet`; Cheng et al., 2025), `WolfPackR` provides a comprehensive framework for studying wolf pack structure and dynamics.

## Functionality

### Installation and usage

The `WolfPackR` package can be installed through the GitHub page (https://github.com/eboncourt/WolfPackR) using the remotes::install_github function. Detailed descriptions of the functions along with examples are available through the help function of `WolfPackR` in the R environment. The package also includes a vignette and a data set example.

WolfPackR was designed with R 4.4.1. The dependencies in the R environment are `igraph` (Csárdi et al., 2025), `dplyr` (Wickham et al., 2026)`, sf` (Pebesma, 2018) and `leaflet` (Cheng et al., 2025).

### Functions

`WolfPackR` provides a comprehensive toolkit of seven functions (Figure 2, Table 1).

**Table 1:** Details of package functions, their inputs, parameters, and outputs.

| Function | Input | Main parameters | Output |
| --- | --- | --- | --- |
| `genetic_groups()` | A data.frame with columns: `'ind1'`, `'ind2'`, and a relatedness estimator. | `'estimator'`: Column name for relatedness estimates. `'threshold'`: Minimum relatedness value for grouping. | A data.frame with individuals assigned to genetic groups. |
| `mcp_sf()` | An `'sf'` object or a data.frame with columns `'X'` and `'Y'`. | `'percentile'`: Percentile for outlier exclusion. `'buffer_radius'`: Buffer for insufficient data points. | An `'sf'` object representing the MCP, or NULL if insufficient points. |
| `calculate_overlap()` | Two polygon objects (`'sf'`). | None | A numeric overlap ratio (0 to 1) between the polygons. |
| `ugly_ducklings()` | An `'sf'` object with individuals and their packs. | `'min_overlap'`: Minimum overlap percentage. `'buffer'`: Buffer distance. `'mcp.percent'`: MCP percentile. | A data.frame classifying individuals as "In Group," "Lone Individual," or "Ugly Duckling." |
| `spatial_groups()` | An `'sf'` object containing observations with group information. | `'percentile'`: MCP percentile. `'buffer_radius'`: Buffer radius. `'max_iterations'`: Max iterations. `'min_mcp_overlap'`: Minimum MCP overlap. | A data.frame with updated spatial subgroup assignments. |
| `summarise_packs()` | An `'sf'` object with individuals and their packs. | `'sex_column'`: Column name for sex information. `'male_pattern'`, `'female_pattern'`: Regex patterns. | A list summarizing pack composition, territory area, and putative dominant individuals. |
| `plot_packs()` | An `'sf'` object with individuals and their packs. | None | An interactive `'leaflet'` map visualizing pack territories and individual locations. |

**Figure 1:**
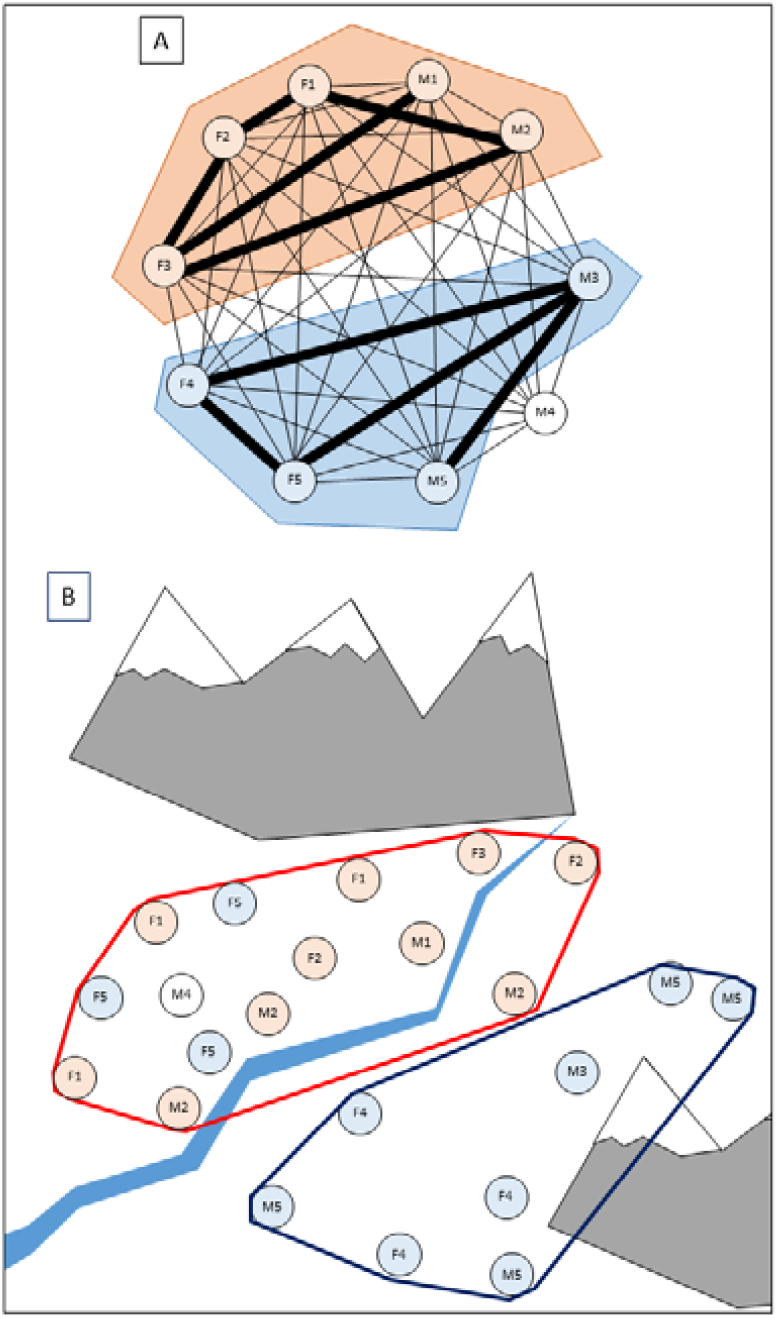
Example illustrating how `WolfPackR` works. A) Based on the pairwise parentage estimators between 10 individuals, a relationship graph is created. The thick lines represent parentage estimators above a threshold chosen by the user. This allows us to determine the existence of three genetic groups of individuals: two represented in colour, and the third consisting solely of individual M4. B) By spatialising the data, we see that individual F5 is not spatially connected to its genetic group, so it is classified as an ‘ugly duckling’. As its observed territory overlaps with that of the other genetic group, it is considered that it and the individuals in this group form a pack (shown in red), while the others are part of a second pack (shown in blue).

**Figure 2:**
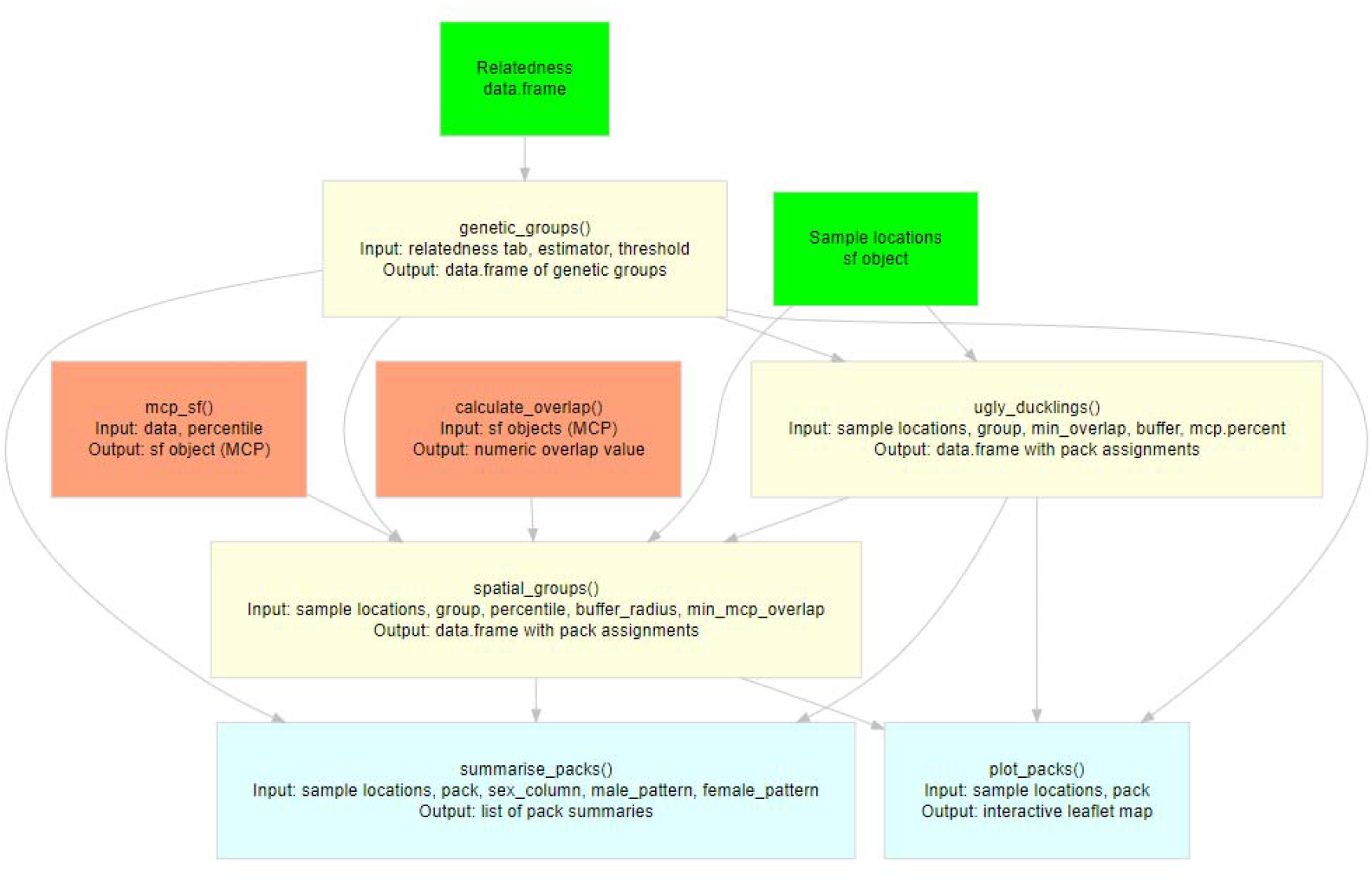
`WolfPackR` functions and possible workflow. Input data are shown in green, main functions in yellow, display functions in blue and auxiliary functions in orange. The typical workflow goes from input data to display, passing through the `genetic_groups`, `ugly_ducklings`, and `spatial_groups` functions, but other combinations are possible to offer users greater flexibility.

These functions collectively enable users to integrate genetic and spatial data, facilitating analyses of population structure, territoriality, and social organization in wildlife studies. The toolkit supports both quantitative summaries and interactive visualizations, enhancing interpretability and reproducibility in ecological research. The modular design allows for flexible application across diverse study systems and research questions.

The `genetic_groups` function is designed to identify genetic groups from a pairwise relatedness matrix using a graph-based clustering approach. This function is particularly useful in population genetics and ecological studies where understanding the genetic structure of populations is crucial. The function takes as input a data frame (`relate`) containing pairwise relatedness estimates between individuals, typically generated by software such as the `related` package (Pew et al., 2015). The function begins by filtering the relatedness matrix to retain only those pairs of individuals with relatedness values exceeding a user-defined threshold. This threshold allows researchers to focus on significant genetic relationships, excluding weaker or spurious connections. The function then constructs an adjacency matrix, which is used to create a graph where nodes represent individuals and edges represent genetic relatedness above the specified threshold. Using the `igraph` package, the function constructs a graph from the adjacency matrix and detects connected components within the graph. Each connected component represents a genetic group, where individuals within the same component are more closely related to each other than to individuals in other components. The function also handles individuals without any significant genetic connections by assigning them to unique groups, ensuring all individuals are included in the output. The output of the `genetic_groups` function is a data frame containing each individual and their assigned genetic group. This output can be used for further analyses, such as spatial mapping of genetic groups or integration with spatial data.

The `ugly_ducklings` function is designed to identify and assign spatially isolated individuals within genetically linked groups to the most suitable social or spatial pack.

The function operates in three main steps:

1. Identification of Lone Individuals: The function first identifies individuals that are spatially isolated from their group. For each individual, it calculates the Minimum Convex Polygon (MCP) of the remaining group members and checks if the individual’s location lies within this MCP. If an individual’s location does not fall within this buffered MCP, it is classified as a “Lone Individual.” This step accounts for individuals with one or two location points, ensuring that even those with limited data are appropriately classified.
2. Identification of Ugly Ducklings: The function then identifies ‘Ugly Ducklings’, which are individuals that are spatially isolated from their group. For each such individual, the function calculates the MCP of the individual and the MCP of the rest of the group. It then computes the overlap ratio between these MCPs. If the overlap ratio is below a specified threshold (`min_overlap`), the individual is classified as an ‘Ugly Duckling’.
3. Reassignment to Suitable Packs: Finally, the function reassigns “Ugly Ducklings” and “Lone Individuals” to the most suitable pack based on spatial overlap. For each isolated individual, the function calculates the spatial overlap between the individual’s MCP and the MCPs of all other groups. The individual is then reassigned to the group with the highest overlap. If no suitable group is found, the individual retains its original genetic group assignment.

The function takes as input an `sf` object (`obs`) containing spatial data for individuals, including their group memberships. The group parameter specifies the column name containing the group identifiers, which can be genetic or spatial groups. The `min_overlap` parameter defines the minimum overlap percentage to consider an individual as integrated into a group, while the buffer parameter specifies the buffer distance around the MCP of the group to consider for inclusion. The `mcp.percent` parameter determines the percentile of points excluded before calculating the MCP. The output of the `ugly_ducklings` function is a data frame containing all individuals, their genetic group, and their assigned pack, including unique identifiers for ‘Lone Individuals’.

The `spatial_groups` function is designed to identify spatial groups or subgroups within predefined groups using Minimum Convex Polygons (MCPs) and clustering based on spatial overlap. It operates through an iterative process to decompose groups into spatially coherent subgroups. Initially, it calculates the MCP for each predefined group or for all individuals if no group information is provided.

The function then identifies spatial subgroups by calculating individual MCPs for each member of the group and constructing an overlap matrix. This matrix indicates the spatial overlap between the MCPs of each pair of individuals. If the overlap between two MCPs exceeds a user-defined threshold (`min_mcp_overlap`), the individuals are considered spatially linked. Using graph theory, the function constructs an adjacency matrix from the overlap matrix and identifies connected components within the graph. Each connected component represents a spatial subgroup, where individuals within the same component are more spatially cohesive with each other than with individuals in other components. The function iterates this process until the subgroups stabilize or until a maximum number of iterations (`max_iterations`) is reached. The function takes as input an `sf` object (`obs`) containing spatial data for individuals, including their coordinates and group memberships. The `group` parameter specifies the column name containing the group identifiers. If no group information is provided, all individuals are treated as a single group. The `percentile` parameter determines the percentile for the MCP calculation, allowing users to exclude a certain percentage of outlier points. The `buffer_radius` parameter specifies a buffer radius to use if the MCP cannot be calculated. The `min_mcp_overlap` parameter defines the minimum spatial overlap required to consider two MCPs as linked. The output of the `spatial_groups` function is a data frame containing each individual and their assigned spatial subgroup. Individuals that cannot be assigned to a subgroup with at least two other individuals are marked as “Lone Individual.”

The `summarise_packs` and `plot_packs` functions provide a way for summarizing and visualizing spatial and social structures within animal populations. The `summarise_packs` function generates detailed summaries of each pack, including the total number of individuals and the number of spatial points recorded for each individual. It also identifies putative dominant individuals, both male and female, based on the number of spatial points recorded, which can serve as a proxy for territory usage or activity levels (Mech, 1999). Additionally, the function calculates the territory area for each pack using MCP, providing a quantitative measure of the spatial extent of each group. The function can infer sex from individual IDs using regex patterns. The `plot_packs` function complements the summary by creating an interactive map that visualizes pack territories and individual locations using the `leaflet` package.

The `mcp_sf` function computes the Minimum Convex Polygon (MCP) for a set of spatial points, with an option to exclude outlier points beyond a specified percentile distance from the centroid. The `calculate_overlap` function quantifies the spatial overlap between two polygons, returning a ratio between 0 and 1. This function is designed to handle various edge cases, including null inputs, empty intersections, and non-polygonal geometries, ensuring robustness across diverse datasets. By providing a standardized measure of spatial overlap, it facilitates the assessment of spatial interactions among individuals or groups.

`WolfPackR` also comes with 4 datasets that allow to explore the different functions. `samples` is an `sf` object containing the coordinates of samples from 9 different fictitious individuals. `relate` is a matrix containing the pairwise estimators of relatedness between these 9 individuals. The other datasets `Bruvo_Carpathians` and `samples_Carpathians` allow you to reproduce the analyses presented below (Case study).

## Case study

### Material and methods

We tested `WolfPackR` on an existing dataset comprising 505 genotyped and geospatialised samples from wolf scats collected in Romania from 2017 to 2020 (Foundation Conservation Carpathia, 2024). This dataset has already been thoroughly analysed to reconstruct individuals’ family trees and use human expertise to delineate pack territories (Iosif et al., 2025). It therefore provides an ideal basis for testing `WolfPackR` and comparing its results with those obtained by other methods.

We started with genotyping data for which we calculated a Bruvo genetic distance (Bruvo et al., 2004) using the R package `polysat` (Clark & Jasieniuk, 2011) between each pair of individuals. We considered similarity to be equal to 1 – Bruvo distance, and set a similarity threshold of 0.7 (i.e. a Bruvo distance of 0.3) to define pairings, in accordance with the genetic structure observed in our dataset, using the `genetic_groups` function. We used several combinations of the different `WolfPackR` functions to illustrate the variety of possible workflows (Figure 3). We considered individuals to be spatially matched on their respective 100% MCPs if they overlapped by more than 50%.

**Figure 3:**
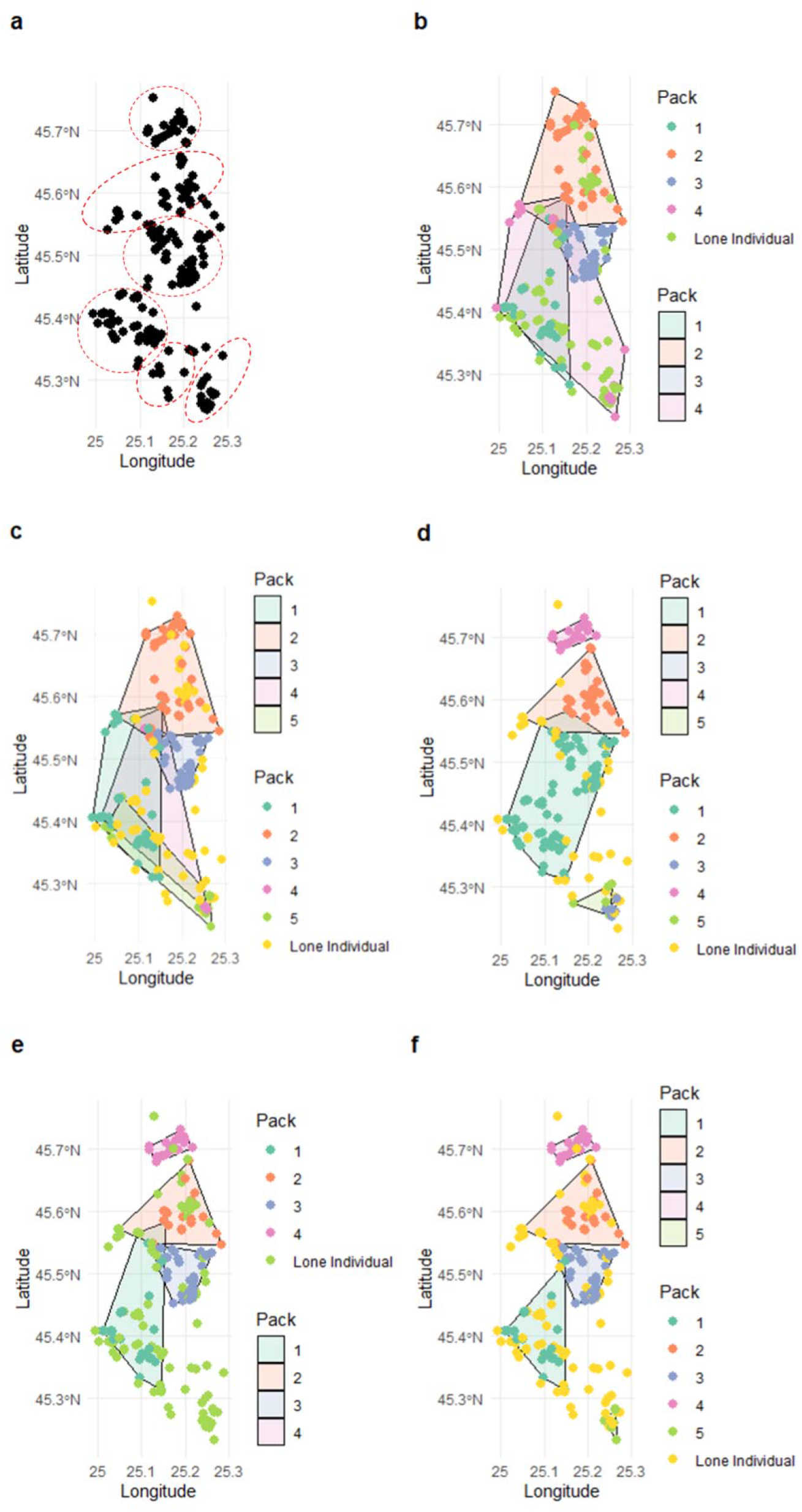
Case studies on a dataset from grey wolf genotyping in Romania (Foundation Conservation Carpathia, 2024). a. Sampling (black dots) and pack territories (red) deduced from the work of Iosif et al. (2025), with human expertise. b. Genetic groups obtained using the `genetic_groups` function with a Bruvo distance. c. Groups obtained using the `genetic_groups` and `ugly_ducklings` functions. d. Groups obtained solely on the basis of spatial data (`spatial_groups` function). e. Packs identified using `genetic_groups` and `spatial_groups`. f. Packs identified using the complete `WolfPackR` workflow: `genetic_groups`, `ugly_ducklings` and `spatial_groups`. In b, c, d, e, and f, each point represents a sample and each polygon represents the MCP of a pack.

## Results

By considering only genetic relationships, four genetic groups can be identified, whose individuals are largely clustered within the study area, which is consistent with the inter-pack dispersal movements described in the analysis by Iosif et al. (2025) (Fig. 3b). This result is refined by the use of `ugly_ducklings` (Fig. 3c). An analysis considering only spatial data, without predefined genetic groups, allows us to roughly identify four spatial groups that resemble the packs identified by Iosif et al. (2025) (Fig. 3a), while seemingly missing some key results (pack 1 appears to be the union of two packs, packs 3 and 5 have small territories that almost overlap) (Fig. 3d). By performing this spatial analysis based on the genetic groups identified with the `genetic_groups` function (Fig. 3e) and, even more so, with the complete workflow incorporating the detection of ugly ducklings (Fig. 3f), the results are closer to those determined by reconstructing family trees and by expert assessment. In particular, five well-defined packs can be distinguished (Fig. 3f), corresponding to five of the six packs identified by Iosif et al. (2025). Only one pack, in the south of the study area, escapes `WolfPackR` due to the low spatial density of data in this location.

## Conclusion

`WolfPackR` enables researchers and practitioners to accurately classify wolf packs using both genetic and spatial data using a reproductive and cost-effective workflow in terms of resources and computing time. It provides functions to visualize pack territories and relationships interactively, all within an R environment allowing for broader integration into an analytical workflow. Its modular design and reliance on widely used R packages (`dplyr`, `igraph`, `sf`, `leaflet`) ensure flexibility and ease of use. Its main limitation lies in the fact that the probability of detecting individuals among grey wolves is highly variable. Insufficient or heterogeneous sampling across the landscape may result in pack territories that do not reflect the biological reality. Future developments may include support for more complex relatedness estimators or the shift to a kernel-based approach rather than an MCP approach for estimating territories. However, the case study presented shows that the package can produce interesting results from a research or management perspective.

## Acknowledgements

We would like to thank Ruben Iosif and Tomaž Skrbinšek for their feedback on the package. We would also like to thank Björn Reineking and Nathan Daumergue for their assistance.

## References

Alexander, S. M., Paquet, P. C., Logan, T. B., & Saher, D. J. (2005). Snow-tracking versus radiotelemetry for predicting wolf-environment relationships in the Rocky Mountains of Canada. Wildlife Society Bulletin, 33(4), 1216□1224. 10.2193/0091-7648(2005)33[1216:SVRFPW]2.0.CO;2

Ausband, D. E., Mitchell, M. S., & Waits, L. P. (2017). Effects of breeder turnover and harvest on group composition and recruitment in a social carnivore. Journal of Animal Ecology, 86(5), 1094□1101. 10.1111/1365-2656.12707

Boitani, L. (2003). Wolf Conservation and Recovery. In L.D. Mech & L. Boitani (Éds.), WolveslZ: Behavior, Ecology, and Conservation (p. 317□340). University of Chicago Press.

Borg, B. L., Brainerd, S. M., Meier, T. J., & Prugh, L. R. (2015). Impacts of breeder loss on social structure, reproduction and population growth in a social canid. Journal of Animal Ecology, 84(1), 177□187. 10.1111/1365-2656.12256

Bruvo, R., Michiels, N. K., D’Souza, T. G., & Schulenburg, H. (2004). A simple method for the calculation of microsatellite genotype distances irrespective of ploidy level. Molecular Ecology, 13(7), 2101□2106. 10.1111/j.1365-294X.2004.02209.x

Calenge, C. (2006). The package “adehabitat” for the R software⍰: A tool for the analysis of space and habitat use by animals. Ecological Modelling, 197(3□4), 516□519. 10.1016/j.ecolmodel.2006.03.017

Caniglia, R., Fabbri, E., Galaverni, M., Milanesi, P., & Randi, E. (2014). Noninvasive sampling and genetic variability, pack structure, and dynamics in an expanding wolf population. Journal of Mammalogy, 95(1), 41□59. 10.1644/13-MAMM-A-039

Chapron, G., Kaczensky, P., Linnell, J. D. C., von Arx, M., Huber, D., Andrén, H., López-Bao, J. V., Adamec, M., žlvares, F., Anders, O., Balčiauskas, L., Balys, V., Bedő, P., Bego, F., Blanco, J. C., Breitenmoser, U., Brøseth, H., Bufka, L., Bunikyte, R., … Boitani, L. (2014). Recovery of large carnivores in Europe’s modern human-dominated landscapes. Science, 346(6216), 1517□1519. 10.1126/science.1257553

Cheng, J., Schloerke, B., Karambelkar, B., Xie, Y., & Aden-Buie, G. (2025). leafletlZ: Create Interactive Web Maps with the JavaScript « Leaflet » Library (Version 2.2.3.9000) [R package]. https://rstudio.github.io/leaflet/

Clark, L. V., & Jasieniuk, M. (2011). POLYSAT⍰: An R package for polyploid microsatellite analysis. Molecular Ecology Resources, 11(3), 562□566. 10.1111/j.1755-0998.2011.02985.x

Csárdi, G., Nepusz, T., Müller, K., Horvát, S., Traag, V., Zanini, F., & Noom, D. (2025). igraph for RlZ: R interface of the igraph library for graph theory and network analysis (Version v2.1.4) [Logiciel]. Zenodo. 10.5281/ZENODO.7682609

Foundation Conservation Carpathia. (2024). Wolf population size and composition in one of Europe’s strongholds, the Romanian Carpathians [Jeu de données]. Zenodo. 10.5281/ZENODO.14544266

Fuller, T. K., Mech, L. D., & Cochrane, J. F. (2003). Wolf Population Dynamics. In L.D. Mech & L. Boitani (Éds.), WolveslZ: Behavior, ecology, and conservation (p. 161□191). University of Chicago Press.

Iosif, R., Skrbinšek, T., Erős, N., Konec, M., Boljte, B., Jan, M., & Promberger-Fürpass, B. (2025). Wolf Population Size and Composition in One of Europe’s Strongholds, the Romanian Carpathians. Ecology and Evolution, 15(4), e71200. 10.1002/ece3.71200

Jones, O. R., & Wang, J. (2010). COLONY⍰: A program for parentage and sibship inference from multilocus genotype data. Molecular Ecology Resources, 10(3), 551□555. 10.1111/j.1755-0998.2009.02787.x

Kelly, M. J., Betsch, J., Wultsch, C., Mesa, B., & Mills, L. S. (2012). Noninvasive sampling for carnivores. In L. Boitani & R.A. Powell (Éds.), Carnivore Ecology and Conservation (p. 47□69). Oxford University Press. 10.1093/acprof:oso/9780199558520.003.0004

López-Bao, J. V., Godinho, R., Pacheco, C., Lema, F. J., García, E., Llaneza, L., Palacios, V., & Jiménez, J. (2018). Toward reliable population estimates of wolves by combining spatial capture-recapture models and non-invasive DNA monitoring. Scientific Reports, 8(1), 2177. 10.1038/s41598-018-20675-9

Manel, S., Schwartz, M. K., Luikart, G., & Taberlet, P. (2003). Landscape genetics⍰: Combining landscape ecology and population genetics. Trends in Ecology & Evolution, 18(4), 189□197. 10.1016/S0169-5347(03)00008-9

Mech, L. D. (1999). Alpha status, dominance, and division of labor in wolf packs. Canadian Journal of Zoology, 77.

Mech, L. D., & Boitani, L. (Éds.). (2003). WolveslZ: Behavior, ecology, and conservation (Repr.). Univ. of Chicago Press.

Pacheco, C., Rio-Maior, H., Nakamura, M., žlvares, F., & Godinho, R. (2024). Relatedness-based mate choice and female philopatry⍰: Inbreeding trends of wolf packs in a human-dominated landscape. Heredity, 132(4), 211□220. 10.1038/s41437-024-00676-3

Pebesma, E. (2018). Simple Features for R⍰: Standardized Support for Spatial Vector Data. The R Journal, 10(1), 439. 10.32614/RJ-2018-009

Pew, J., Muir, P. H., Wang, J., & Frasier, T. R. (2015). related⍰: An R package for analysing pairwise relatedness from codominant molecular markers. Molecular Ecology Resources, 15(3), 557□561. 10.1111/1755-0998.12323

Pilot, M., Moura, A. E., Okhlopkov, I. M., Mamaev, N. V., Manaseryan, N. H., Hayrapetyan, V., Kopaliani, N., Tsingarska, E., Alagaili, A. N., Mohammed, O. B., Ostrander, E. A., & Bogdanowicz, W. (2021). Human-modified canids in human-modified landscapes⍰: The evolutionary consequences of hybridization for grey wolves and free-ranging domestic dogs. Evolutionary Applications, 14(10), 2433□2456. 10.1111/eva.13257

Pritchard, J. K., Stephens, M., & Donnelly, P. (2000). Inference of Population Structure Using Multilocus Genotype Data. Genetics, 155(2), 945□959. 10.1093/genetics/155.2.945

Redpath, S. M., Linnell, J. D. C., Festa-Bianchet, M., Boitani, L., Bunnefeld, N., Dickman, A., Gutiérrez, R. J., Irvine, R. J., Johansson, M., Majić, A., McMahon, B. J., Pooley, S., Sandström, C., Sjölander-Lindqvist, A., Skogen, K., Swenson, J. E., Trouwborst, A., Young, J., & Milner-Gulland, E. J. (2017). Don’t forget to look down – collaborative approaches to predator conservation. Biological Reviews, 92(4), 2157□2163. 10.1111/brv.12326

Royle, J. A., Chandler, R. B., Sollmann, R., & Gardner, B. (Éds.). (2014). Spatial capture-recapture. Elsevier.

Rutledge, L. Y., Patterson, B. R., Mills, K. J., Loveless, K. M., Murray, D. L., & White, B. N. (2010). Protection from harvesting restores the natural social structure of eastern wolf packs. Biological Conservation, 143(2), 332□339. 10.1016/j.biocon.2009.10.017

Schwartz, M., Luikart, G., & Waples, R. (2007). Genetic monitoring as a promising tool for conservation and management. Trends in Ecology & Evolution, 22(1), 25□33. 10.1016/j.tree.2006.08.009

Taberlet, P. (1996). Reliable genotyping of samples with very low DNA quantities using PCR. Nucleic Acids Research, 24(16), 3189□3194. 10.1093/nar/24.16.3189

Treves, A., & Karanth, K. U. (2003). Human-Carnivore Conflict and Perspectives on Carnivore Management Worldwide. Conservation Biology, 17(6), 1491□1499. 10.1111/j.1523-1739.2003.00059.x

Wang, J. (2002). An Estimator for Pairwise Relatedness Using Molecular Markers. Genetics, 160(3), 1203□1215. 10.1093/genetics/160.3.1203

Wickham, H., François, R., Henry, L., Müller, K., & Vaughan, D. (2026). dplyrlZ: A Grammar of Data Manipulation (Version 1.2.0) [R package]. https://dplyr.tidyverse.org

